# Chromosome assembly for the meagre, *Argyrosomus regius*, reveals species adaptations and sciaenid sex-related locus evolution

**DOI:** 10.1101/2022.10.04.510760

**Authors:** Vasileios Papadogiannis, Tereza Manousaki, Orestis Nousias, Alexandros Tsakogiannis, Jon B. Kristoffersen, Constantinos C. Mylonas, Constantinos Batargias, Dimitrios Chatziplis, Costas S. Tsigenopoulos

## Abstract

The meagre, *Argyrosomus regius*, has recently become a species of increasing economic interest for the Mediterranean aquaculture and there is ongoing work to boost production efficiency through selective breeding. Access to the complete genomic sequence will provide an essential resource for studying quantitative trait-associated loci and exploring the genetic diversity of different wild populations and aquaculture stocks in more detail. Here, we present the first complete genome for *A. regius*, produced through a combination of long and short read technologies and an efficient in-house developed pipeline for assembly and polishing. Scaffolding using previous linkage map data allowed us to reconstruct a chromosome level assembly with high completeness, complemented with gene annotation and repeat masking. We use this new resource to study the evolution of the meagre genome and other Sciaenids, via a comparative analysis of 25 high-quality teleost genomes. Combining a rigorous investigation of gene duplications with base-wise conservation analysis, we identify candidate loci related to immune, fat metabolism and growth adaptations in the meagre. Following phylogenomic reconstruction, we show highly conserved synteny within Sciaenidae. In contrast, we report rapidly evolving syntenic rearrangements and gene copy changes in the sex-related *dmrt1* neighbourhood in meagre and other members of the family. These novel genomic datasets and findings will add important new tools for aquaculture studies and greatly facilitate husbandry and breeding work in the species.

## Introduction

The meagre, *Argyrosomus regius*, is a teleost fish in the family Sciaenidae, commonly known as croakers or drums because of the characteristic croaking sounds they produce. Sciaenids include some commercially important species with sequenced genomes, such as the related Japanese meagre *(Argyrosomus japonicus*)(Zhao et al., 2021), the large yellow croaker (*Larimichthys crocea*)(Ao et al., 2015), the red drum (*Sciaenops ocellatus*)(T. Xu, Li, Chu, & Zheng, 2021) and the spinyhead croaker (*Collichthys lucidus*)(Cai et al., 2019).

Several favourable characteristics of the species, such as fast growth, large body size and low-fat content have attracted interest in meagre aquaculture in recent years, promoting efforts to improve hatchery techniques and culture efficiency(Mylonas, Mitrizakis, Papadaki, & Sigelaki, 2013). Global meagre production amounted to more than 55,500 tonnes in 2019, with 68% of this volume sourced from aquaculture, while the European Union was the second largest producer in the world after Egypt (www.fao.org & www.apromar.es). Previous work on meagre aquaculture has looked into the application of varying dietary sources for culturing the species(Chatzifotis, Panagiotidou, & Divanach, 2012; Chatzifotis et al., 2010), including vegetable(Ribeiro et al., 2015) and insect-based diets(Guerreiro et al., 2020), the study of the development of the digestive system(Papadakis, Kentouri, Divanach, & Mylonas, 2013), as well as meagre reproduction and spawning(Mylonas, Mitrizakis, Castaldo, et al., 2013; Mylonas, Mitrizakis, Papadaki, et al., 2013; Ramos-Júdez et al., 2019).

The recent increased interest in improving meagre breeding and culture has created the need for genetic resources for the species. In this direction, there has been prior research on the development of genetic markers for the study of growth traits, including the meagre muscle and liver transcriptomes(Manousaki et al., 2018), a microsatellite PCR panel(Vallecillos et al., 2022) a broodstock structure analysis(Orestis Nousias et al., 2021) and a recent meagre ddRAD linkage map(O. Nousias et al., 2022). While such efforts provide valuable tools for meagre aquaculture, they remain constrained in the absence of a complete nuclear genomic sequence and would greatly benefit from the availability of a high-quality assembly for the species.

In this study, we present the first nuclear genome for the meagre. Using a combination of long and short read data and the previously published linkage map, we built a chromosome level assembly with high completeness and provide high quality repeat and gene annotations. Using the new assembly, we carried out phylogenetic and evolutionary analyses, focusing on gene duplications and signatures of accelerated evolution in meagre genes. An investigation of duplicated families and fast evolving loci highlighted immune-system, fat metabolism and cancer-related adaptations, paving the way for understanding the rapid growth and large body size of the species. Finally, exploring of synteny around the sex-related *dmrt1* locus in sciaenids revealed rapid changes in the genomic neighbourhood in the meagre, offering a primary candidate for follow up reproductive studies in the species.

## Materials and Methods

### Sampling and Sequencing

Genomic material for sequencing was isolated from a female *A. regius* individual, collecting a total of 10ml blood in 1/10 volume of heparin. DNA for long read sequencing on an Oxford Nanopore MinION sequencer was extracted on the same day of the collection of 2 ml blood, using the Qiagen Genomic tip (20 G), following the Oxford Nanopore Technologies protocol for DNA extraction from chicken blood. Assessment of DNA integrity quality was done through electrophoresis on a 0.4% w/v Bio-Rad Megabase agarose gel. DNA purity was estimated by Nanodrop ratios and DNA concentration by Qubit. Library preparation was then carried out following manufacturer instructions, using the sequencing kit SQK-LSK109. Two libraries were sequenced for 96 hours on two R9.4.1 flow cells on the MinION sequencer of IMBBC, HCMR. Raw reads were basecalled with Guppy 4.0.11, using the High Accuracy (hac) configuration. A total of 3,003,301 reads were generated with a quality score above 7, with an N50 of 30,600, totalling 37,962,807,176 bp and 99.94% passed quality control (Supplementary Table 1). For Illumina sequencing, two-day old frozen blood was used, following the same extraction protocol described above, similar to Danis et al. 2021(Danis et al., 2021). Extracted DNA was fragmented via sonication, followed by PCR free library preparation with the Kapa Hyper Prep DNA kit and paired end (150bp) sequencing on an Illumina Hiseq4000 platform (Norwegian Sequencing Center, NSC). A total of 143,224,530 150bp reads were generated, with 85.81% passing quality control (Supplementary Table 1).

RNA extraction was done from material for tissues shown in table 1, homogenised in TRIzol reagent (Invitrogen), under liquid nitrogen. Whole RNA was extracted from the homogenised material following manufacturer instructions. Library preparation for paired end sequencing (150bp) was done with the Illumina TruSeqTM RNA Sample Preparation Kit v2 and libraries were sequenced on the above Illumina Hiseq4000 platform (NSC). More than 100 million 150bp reads were generated for each tissue and more than 90% of reads passed quality control for any tissue (Supplementary Table 2).

**Table 1.**
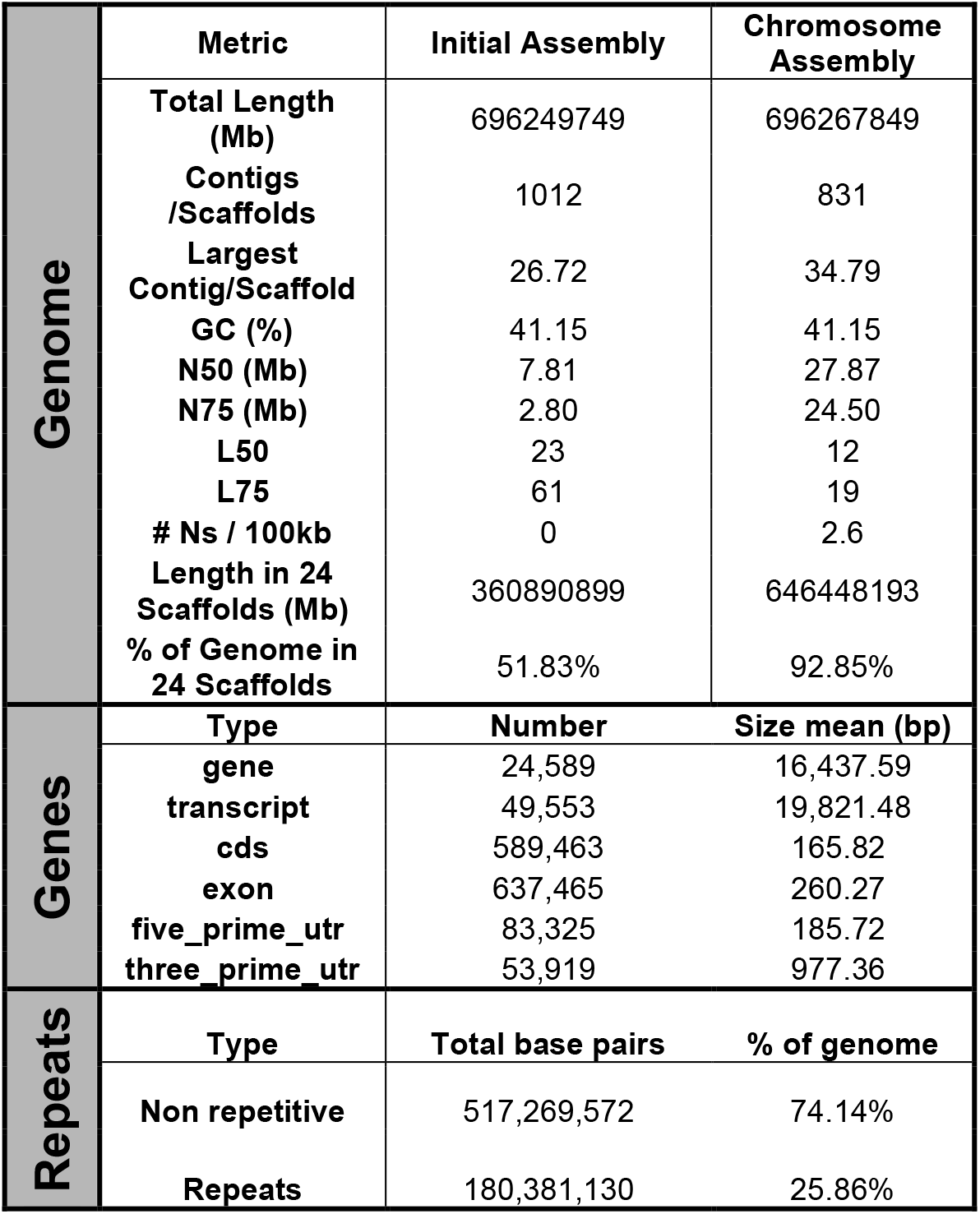
Assembly and annotation metrics.

| Genome | Metric | Initial Assembly | Chromosome Assembly |
| --- | --- | --- | --- |
|  | Total Length (Mb) | 696249749 | 696267849 |
|  | Contigs /Scaffolds | 1012 | 831 |
|  | Largest Contig/Scaffold | 26.72 | 34.79 |
|  | GC (%) | 41.15 | 41.15 |
|  | N50 (Mb) | 7.81 | 27.87 |
|  | N75 (Mb) | 2.80 | 24.50 |
|  | L50 | 23 | 12 |
|  | L75 | 61 | 19 |
|  | # Ns / 100kb | 0 | 2.6 |
|  | Length in 24 Scaffolds (Mb) | 360890899 | 646448193 |
|  | % of Genome in 24 Scaffolds | 51.83% | 92.85% |
| Genes | Type | Number | Size mean (bp) |
|  | gene | 24,589 | 16,437.59 |
|  | transcript | 49,553 | 19,821.48 |
|  | cds | 589,463 | 165.82 |
|  | exon | 637,465 | 260.27 |
|  | five_prime_utr | 83,325 | 185.72 |
|  | three_prime_utr | 53,919 | 977.36 |
| Repeats | Type | Total base pairs | % of genome |
|  | Non repetitive | 517,269,572 | 74.14% |
|  | Repeats | 180,381,130 | 25.86% |

### Assembly and Scaffolding

A kmer counting/distribution strategy was used for genomic data quality assessment, using jellyfish (v2.3.0) for kmer counting (21 bp length) and genomescope (v1.0) to calculate kmer distribution plots(Vurture et al., 2017).

Assembly construction was carried out with the SnakeCube containerised pipeline(Angelova, Danis, Lagnel, & Tsigenopoulos, n.d.). Briefly, sequencing data quality control and trimming is carried out via Trimmomatic(Bolger, Lohse, & Usadel, 2014) (v0.39) and fastQC(*FastQC*, 2015) (v0.11.8) for short read data and via Nanoplot(de Coster, D’Hert, Schultz, Cruts, & van Broeckhoven, 2018) (1.29.0) and porechop (v0.2.3) (https://github.com/rrwick/Porechop#license) for long read data. An initial long read based assembly is built using Flye(Kolmogorov, Yuan, Lin, & Pevzner, 2019) (v2.6), followed by long read based polishing through Racon (v1.4.12) (https://github.com/isovic/racon) and Medaka (v0.9.2) (https://github.com/nanoporetech/medaka). Pilon(Walker et al., 2014) (v1.23) is then used for error correction and polishing using the short read data. Assembly quality, contiguity and completeness were assessed via Quast(Gurevich, Saveliev, Vyahhi, & Tesler, 2013) (v5.0.2), Busco(Simão, Waterhouse, Ioannidis, Kriventseva, & Zdobnov, 2015) (v5.1.0)) and Merqury(Rhie, Walenz, Koren, & Phillippy, 2020).

ALLMAPS(Tang et al., 2015) was used for the scaffolding of assembly contigs based on the meagre Double digest restriction-site associated DNA (ddRAD) linkage map by Nousias et al. (2022) and the species diploid number of chromosomes (2n=48, Soares et al., 2012(Soares et al., 2012)). Preliminary scaffolding efforts suggested that the previous linkage group (LG) I was an artificial merge of two constituent groups, based on non-overlapping mapping of assembly contigs and the unexpectedly large linkage group size compared to the next largest group. Additionally, poor mapping for LG XXIV and its small content in genetic markers suggested that the latter LG was an artifact. After discarding LG XXIV and splitting LG I, we obtained the final scaffolding results presented in this study.

### Repeat Element Annotation

RepeatModeller(Smit, 2015) (v10.1) was used for *de novo* repeat modelling with the Repbase database(Bao, Kojima, & Kohany, 2015), followed by repeat identification and annotation via RepeatMasker(Smit, 2015) (v 4.1.2-p1) while ltr_finder(Z. Xu & Wang, 2007) (v1.07) was used for additional LTR retrotransposon annotation. Repeats identified from these tools were merged to a final repeat annotation set via RepeatCraft(Wong & Simakov, 2019).

### Transcriptome Assembly and Gene Annotation

Transcriptome data quality control were performed with fastQC and trimming with trimmomatic. Reads were mapped to the genome with STAR(Dobin et al., 2013) (v2.7.9a) and separate transcriptome assemblies were built for each tissue via StringTie(Kovaka et al., 2019) (v2.1.4). These transcriptomes were then merged to a final consensus transcriptome assembly via TACO(Niknafs, Pandian, Iyer, Chinnaiyan, & Iyer, 2017).

Based on the transcriptome data, transcript model prediction was done via Mikado(Venturini, Caim, Kaithakottil, Mapleson, & Swarbreck, 2018) (v2.0), incorporating information from intron-exon junction prediction carried out through portcullis (v1.2.0) (https://github.com/EI-CoreBioinformatics/portcullis), open reading frame identification with TransDecoder (v5.5.0) (https://github.com/TransDecoder/TransDecoder) and homology evidence by searching teleost fish and other vertebrate proteomes (dataset presented in Supplementary Table 3) with DIAMOND(Buchfink, Reuter, & Drost, 2021) (v0.9.14). Mikado produced transcript sets were used for training and de-novo gene model prediction with Augustus(Stanke & Morgenstern, 2005) (v3.4.0). Mikado and Augustus datasets were then merged to a final gene annotation set using PASA(Haas, 2003) (v2.4.1). BUSCO was used to assess the completeness of gene annotation datasets. Functional annotation of gene models was carried out through a combination of annotations from PANTHER(Mi & Thomas, 2009) (v2.0) and EggNOG(Huerta-Cepas et al., 2019) (v5.0).

### Orthogroup Identification and Phylogenetic Analyses

OrthoFinder(Emms & Kelly, 2015, 2019) (v2.5.2) was used for gene family inference and one-to-one ortholog identification for phylogenetic reconstruction. After species tree inference, the reconstructed phylogeny was used to re-run the OrthoFinder analysis with the new species tree, with final orthogroups obtained from this corrected run used for downstream analyses.

Alignments of single copy orthologous proteins were built using MAFFT(Katoh, 2002) (v7.4.80), followed by alignment trimming with trimAl(Capella-Gutierrez, Silla-Martinez, & Gabaldon, 2009) (strict mode) (v1.4.rev15). Trimmed alignments were then concatenated to a final super-alignment (super-matrix), which was used for species phylogenetic tree inference through maximum likelihood analysis with RAxML-ng(a Stamatakis, Ludwig, & Meier, 2003; A. Stamatakis, 2014) (v1.0.2), using the JTT model selected by ModelTest-NG(Darriba et al., 2020) and bootstrap resampling with 1,000 bootstrap replicates.

### Synteny Analysis

Macro-synteny analysis in Sciaenidae was based on single copy orthologous loci shared by the meagre and all other species, previously identified from the OrthoFinder analysis. Whole genome synteny plots for single copy orthologs were plotted using Circos(Krzywinski et al., 2009) (v0.69-8).

Synteny in the *dmrt1* neighbourhood was searched using publicly available NCBI data or BLAST(Camacho et al., 2009) (v2.11.0) to investigate if genes of interest are proximal in species without annotated loci in NCBI. For this purpose, we used an *e-value* threshold of 10^−6^ and a similarity threshold of 70% to assess if matches to protein queries were co-located in the same genomic scaffold in each species, recording the genomic location of the genes.

### Gene Duplication Identification and Analysis

To identify potential gene duplication events, we combined two tools that use different approaches; CAFE (v4)(M. v. Han, Thomas, Lugo-Martinez, & Hahn, 2013) calculates rapid gene family expansions and contractions based on gene count tables in each family in each species, while GeneRax(Morel, Kozlov, Stamatakis, & Szoll}osi, 2020) (v2.0.2) carries out gene tree reconciliation based on the species phylogeny and infers putative duplication events from the corrected gene trees. For CAFE, we used gene count matrices produced for OrthoFinder orthogroups, with a *p-value* threshold of 0.01, after discarding orthogroups with low representation across our phylogeny (genes present in less than 4 species) or with no representation in the meagre. For GeneRax input, we built alignments (MAFFT), starting trees (IQ-TREE(Nguyen, Schmidt, von Haeseler, & Minh, 2015) v1.6.12) and calculated phylogenetic models (IQ-TREE), using our reconstructed phylogeny and the UndatedDL model (default mode). We then filtered the output of both tools, keeping the orthogroups suggested as containing duplications by both tools that had two or more duplications in CAFE.

### Multiple Whole-Genome Alignment and Conservation Score Analyses

Based on our reconstructed phylogeny, multiple whole genome alignments for all 25 genomes included in this study were carried out via CACTUS(Armstrong et al., 2020) (v 2.2.1). Based on a multiple genome alignment for each meagre chromosome sequence and our reconstructed phylogeny, we used phyloFit(Siepel & Haussler, 2004) for phylogenetic model fitting for each chromosomal alignment (using the REV substitution model) and phyloP(Pollard, Hubisz, Rosenbloom, & Siepel, 2010) to calculate base-wise conservation scores (using the SPH method and CONACC mode for p-value computation) for each chromosome. Average phyloP (CONACC) scores were calculated for 50kb windows for the 24 meagre chromosomes and plotted using Circos, while average conservation scores were also calculated for all genes and transcripts.

### Gene Ontology Enrichment

Ontology enrichment was carried out via gProfiler(Raudvere et al., 2019). Ontology enrichment for fast evolving genes (with phyloP score<0) was carried out using the “ordered list” option, ranking genes by decreasing conservation score, using our meagre functional annotation reference set. To carry out ontology enrichment for OrthoFinder orthogroups with gene duplications, we built an orthogroup annotation reference set, by leveraging non-redundant ontology annotations from meagre and zebrafish genes belonging to each family. Orthology enrichment for duplication containing orthogroups was then carried out as ordered lists, ranked by number of duplications for each species in the phylogeny. Enriched terms for duplications retrieved for each species from this pipeline were then filtered and ranked by adjusted *p*-values (0.01 threshold). These ranked lists were then compared, retaining only meagre terms that were enriched in a maximum of three other species, to discard terms commonly enriched in duplications across species and shortlist terms that are more likely to be specific to meagre duplications.

### Computational Resources

All computational work described for genome assembly, gene and repeat element annotation, phylogenomic analyses, multiple whole genome alignment and downstream evolutionary analyses were carried out using the computational resources of the IMBBC HPC facility “Zorbas” of HCMR(Zafeiropoulos et al., 2021).

### Ethics and Permits

This study does not include special animal treatment, only euthanasia and study of the genetic material. Thus, the work carried out in the scope of the study does not fall within the HCMR code for animal ethics. Furthermore, we followed all appropriate guidelines for animal care and handling (Guidelines for the treatment of animals in behavioral research and teaching. Anim. Behav. 53, 229–234 (1997))

## Results

### Assembly and scaffolding

For the construction of the genomic assembly of *A. regius*, we used a combination of ONT minion long read data (>54X coverage) and Illumina short read data (>30X coverage). An initial assessment of sequencing quality of the selected specimen was performed using the short read data, revealing very low levels of heterozygosity (predicted range 0.241%-0.244% with average of 0.242%,) and sequencing error (orange line in Fig. 1a). Assembly of the genome was carried out using the SnakeCube LSGA pipeline and this initial assembly was then scaffolded using the high-density ddRAD based linkage map for the species by Nousias et al. (2022). This led to merging 205 contigs into 24 scaffolds corresponding to the species chromosome number (Figure 1b). The final chromosome level assembly has 696.267 Mb of sequence in 831 contigs (N50 = 27.87 Mb, L50 = 12), with 92.85% of the total length contained in the 24 chromosomal scaffolds (Table 1). Assembly completeness was calculated at 96.915% when assessed through kmer counting using the short read data via Merqury (Fig.1c) and at 98.7% when assessed through BUSCO analysis using the Actinopterygii v10 conserved single copy ortholog database (Fig. 1d).

**Figure 1.**
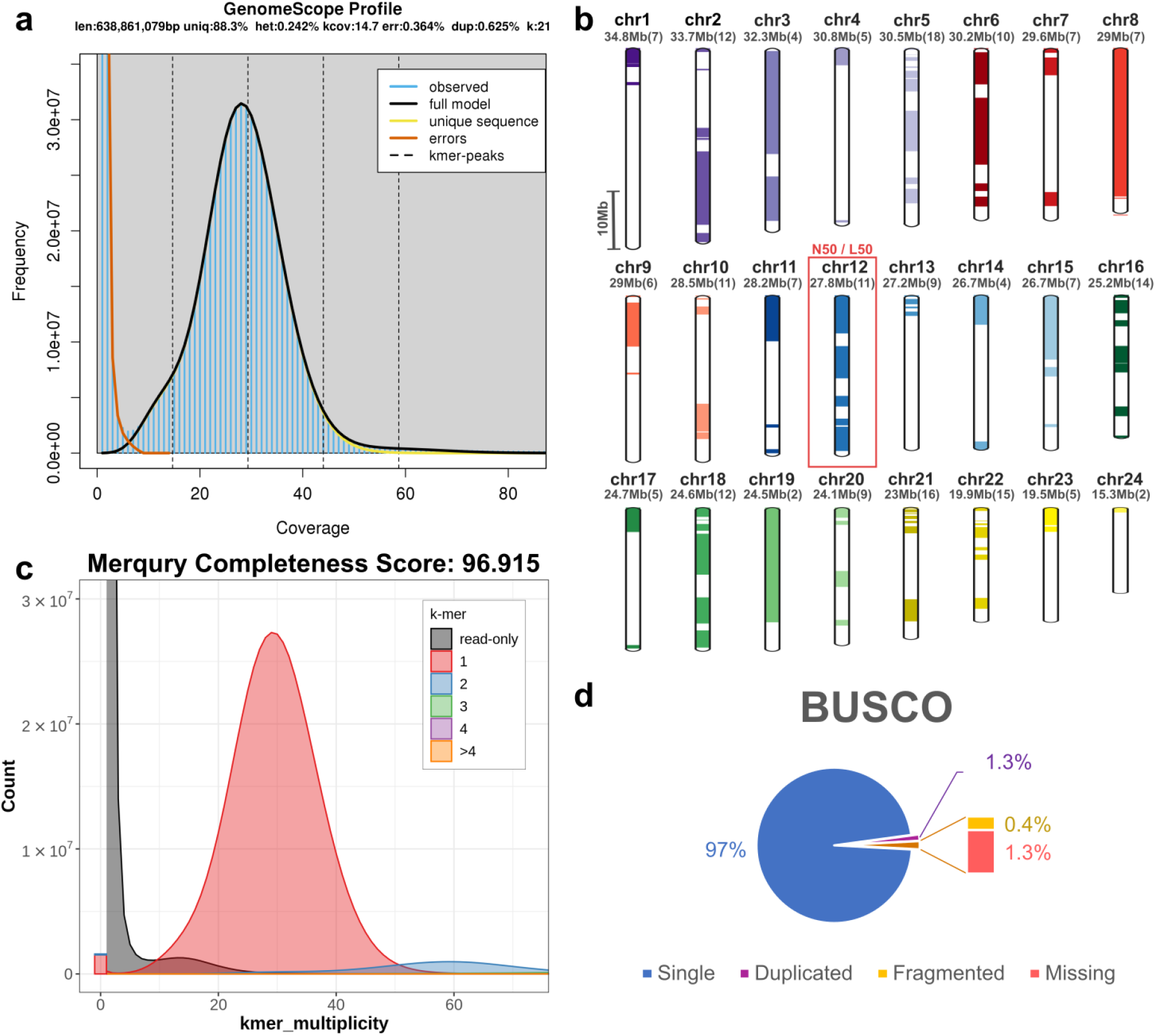
Sequencing and assembly quality assessment. **a**. GenomeScope kmer distribution (kmer length = 21) for short reads used for genome assembly. **b**. Scaffolding of 205 contigs (coloured and white bars) into 24 chromosomal scaffolds, based on the ddRAD linkage map. The red box marking Chromosome 12 corresponds to the N50 and L50 of the assembly. **c**. Merqury kmer distributions (kmer length = 21) for meagre genome assembly, plotting kmers found in 1 to 4 or more copies in the assembly, or only in reads separately. **d**. BUSCO analysis results identifying 98.7% of ultra-conserved single copy orthologs from the actinopterygii10 database, with 1.3% and 0.4% duplicated and fragmented BUSCO orthologs respectively.

### Structural and functional annotation

To identify repeat elements, we used RepeatModeller to obtain species specific repeats models, RepeatMasker and ltr_finder to identify different repeat classes and RepeatCraft to merge all other sets to a final repeat annotation set. The total repeat content of the genome was calculated at 25.86% of the total length, while the genome wide distribution of repeat elements is presented in Figure 2.

**Figure 2.**
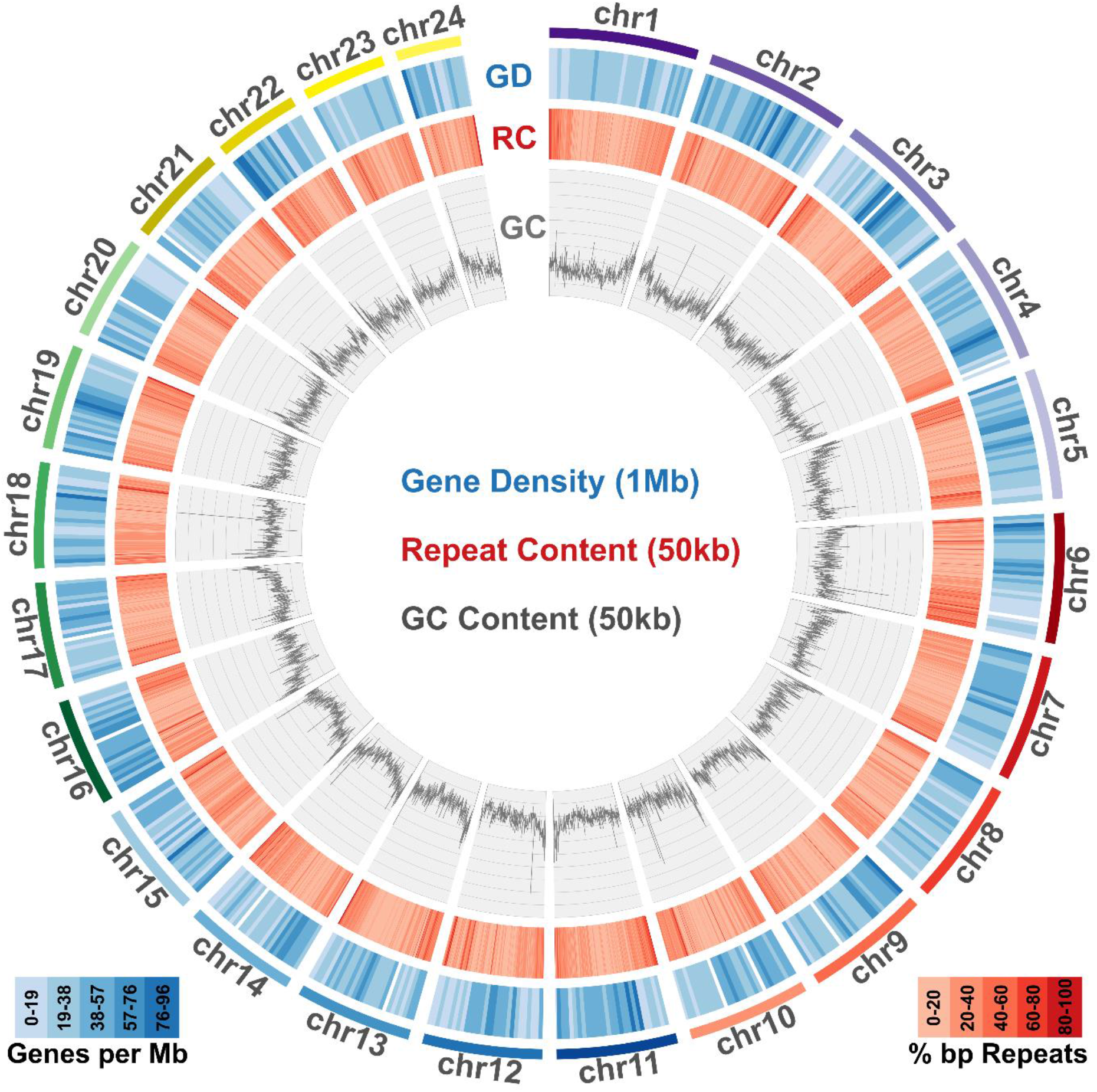
Genome content in genes, repeats and GC. Meagre chromosomal scaffolds are plotted as coloured bars in the outer circle, followed by the gene density heatmap (over 1Mb windows) in the blue circle, the repeat content heatmap (over 50kb windows) in the red circle and the GC content distribution plot (over 50kb windows) in the innermost circle.

Transcriptome data acquired from eight tissues via Illumina sequencing was used to guide gene prediction and annotation. Transcriptome models built using Mikado were provided as a training dataset for Augustus, which was used for *de novo* gene prediction. The acquired de novo gene models were then merged with the Mikado transcript models through the PASA pipeline. The final gene set comprised a total of 24,589 loci, with 637,465 exons and 49,553 transcripts (Table 1). Gene density distribution across the genome and global GC content over a 50kb sliding window are presented in Figure 2.

### Phylogenomic and Gene Duplication analysis of the meagre genome

Taking advantage of the high quality and high completeness of the genome, we used the predicted gene models to characterize meagre gene homology to other teleost species. Based on the assigned orthogroups, we obtained 2,104 single copy orthologous loci present across teleosts (present in all 25 species included), which we used to carry out phylogenomic reconstruction for the 25 species in our study. This phylogeny confidently resolved the expected placement of the meagre in Sciaenidae (all branches obtained with 100 bootstrap support), with the European sea bass (*Dicentrarchus labrax*) placed as an outgroup to the family (Figure 3a). Based on these single copy loci, meagre proteins appear to be evolving comparatively slower compared to other members of the family and most teleosts, based on the short branch length of *A. regius* in the tree. Using divergence time estimates from TimeTree(Kumar et al., 2022), we also calculated divergence times for the meagre at 22.4 million years ago from other members of the family and Sciaenidae as a group at 79.3 million years ago from the European sea bass.

**Figure 3.**
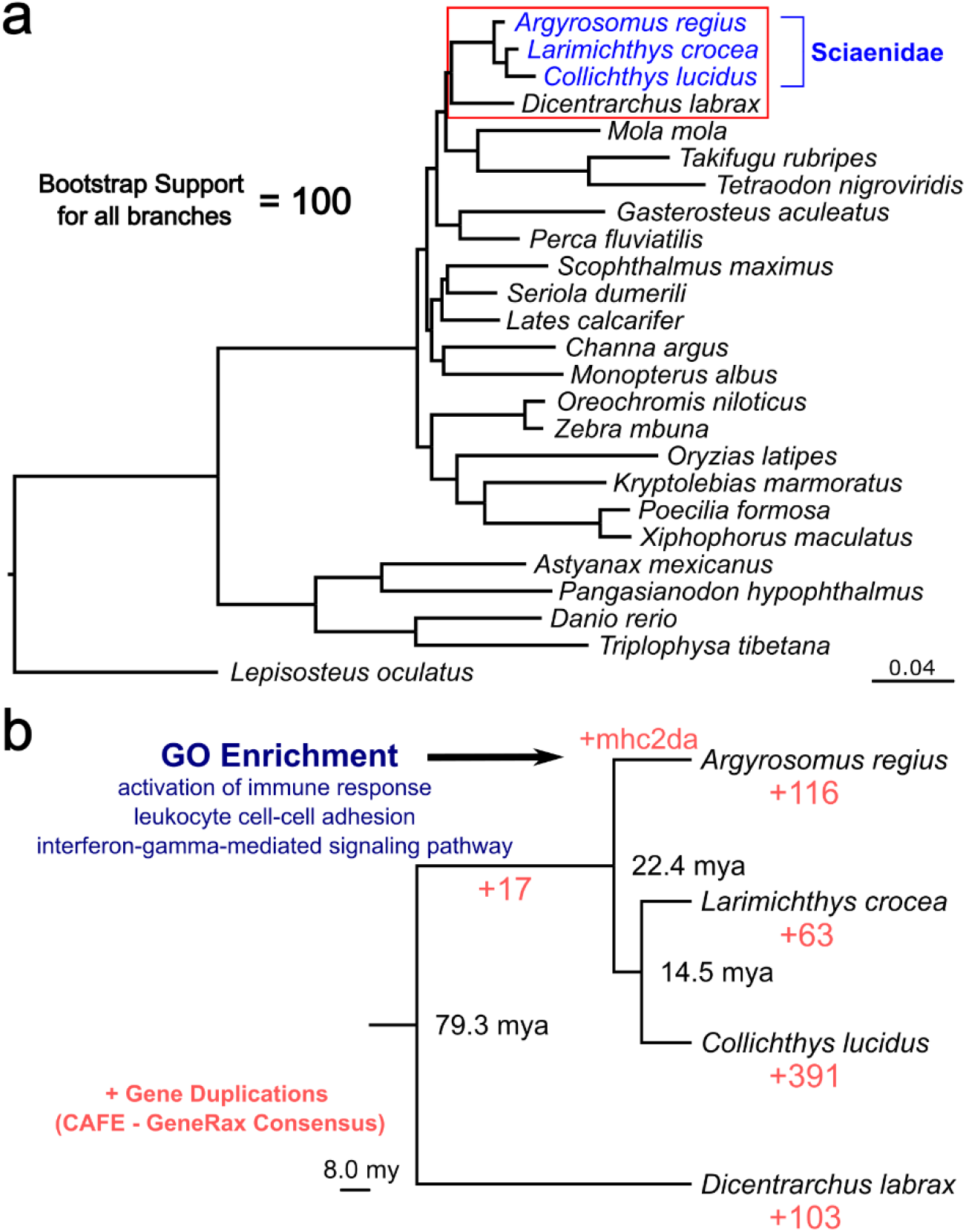
Phylogenomic placement of the meagre and gene duplications in Sciaenidae. **a**. Reconstruction of the evolutionary phylogenetic relationship of 25 fish species, based on 2,104 single copy orthologous loci, placing the meagre in the Sciaenidae family (blue). Bootstrap support for all branches was calculated at 100%. **b**. Focused ultra-metric tree of Sciaenidae and the European sea bass as the outgroup (clade in red box in Fig.3a), with predicted divergence times for each branch shown at branch nodes. Red numbers designate total number of orthogroups with gene duplication events. Selected top terms identified from gene ontology enrichment analysis of meagre duplications are shown in blue.

Combining homology and phylogenomic information, we then characterised gene duplication events in the meagre and compared it to other Sciaenidae and teleosts. Filtering duplication predictions from the two complementary approaches described in materials and methods, we shortlisted 116 orthogroups with duplications in the meagre and 17 orthogroups with duplications in the Sciaenid ancestor. Using the filtered duplication datasets from all species of our phylogeny, we performed comparative ranked (by number of duplications) gene ontology enrichment analysis through gProfiler, to isolate functions associated specifically with meagre duplications. For this purpose, we kept ontology terms significantly enriched in the meagre that are significantly enriched in a maximum of three other species. This led to the shortlisting of 32 ontology terms, mainly related to immune system regulation, immune-related signalling pathway activation and leukocyte cell-cell adhesion (Fig3.b, Table 2). A notable expansion in MHC2 genes was also included in the orthogroups associated with these functions.

**Table 2.** Gene Duplications Ontology Enrichment. Selected top terms from GO enrichment analysis of meagre duplications. Certain terms were omitted to avoid redundancy, with full table available in Supplementary Table 6.

| term_id | term_name | adjusted_p_value |
| --- | --- | --- |
| GO:0002253 | activation of immune response | 1.00E-04 |
| GO:0007159 | leukocyte cell-cell adhesion | 3.88E-04 |
| GO:0002757 | immune response-activating signal transduction | 3.43E-03 |
| GO:0060333 | interferon-gamma-mediated signaling pathway | 5.88E-03 |
| GO:0031347 | regulation of defense response | 8.02E-03 |
| GO:0098742 | cell-cell adhesion via plasma-membrane adhesion molecules | 8.48E-03 |
| GO:0002404 | antigen sampling in mucosal-associated lymphoid tissue | 9.47E-03 |
| GO:2000258 | negative regulation of protein activation cascade | 9.48E-03 |
| GO:0002218 | activation of innate immune response | 1.22E-02 |
| GO:0050870 | positive regulation of T cell activation | 1.38E-02 |

### Accelerated evolution in meagre gene duplications

Complementing gene duplication analyses, we searched for signatures of selection across the meagre genome, after multiple whole genome alignment of the 25 genomes included in the study. Through this analysis, 15,206 transcripts from 8,368 genes were characterised as potentially fast evolving (phyloP CONACC score <0) and 32,379 transcripts from 15,299 genes were characterised as slow evolving (phyloP CONACC score >0). In Figure 4a, we present average conservation scores for the top 10% fastest (836 genes with average phyloP score <=-0.195) and slowest (1529 genes with average phyloP score >= 0.317) evolving loci. Across the meagre genome, chromosome 22 has the highest total number of the top 10% fastest loci (85 genes; 10.1% of fastest genes) and the highest ratio relative to its size (4.27 genes/Mb), followed by chromosome 3 with the second highest total (82 genes; 9.8% of fastest genes) and ratio (2.53 genes/Mb).

**Figure 4.**
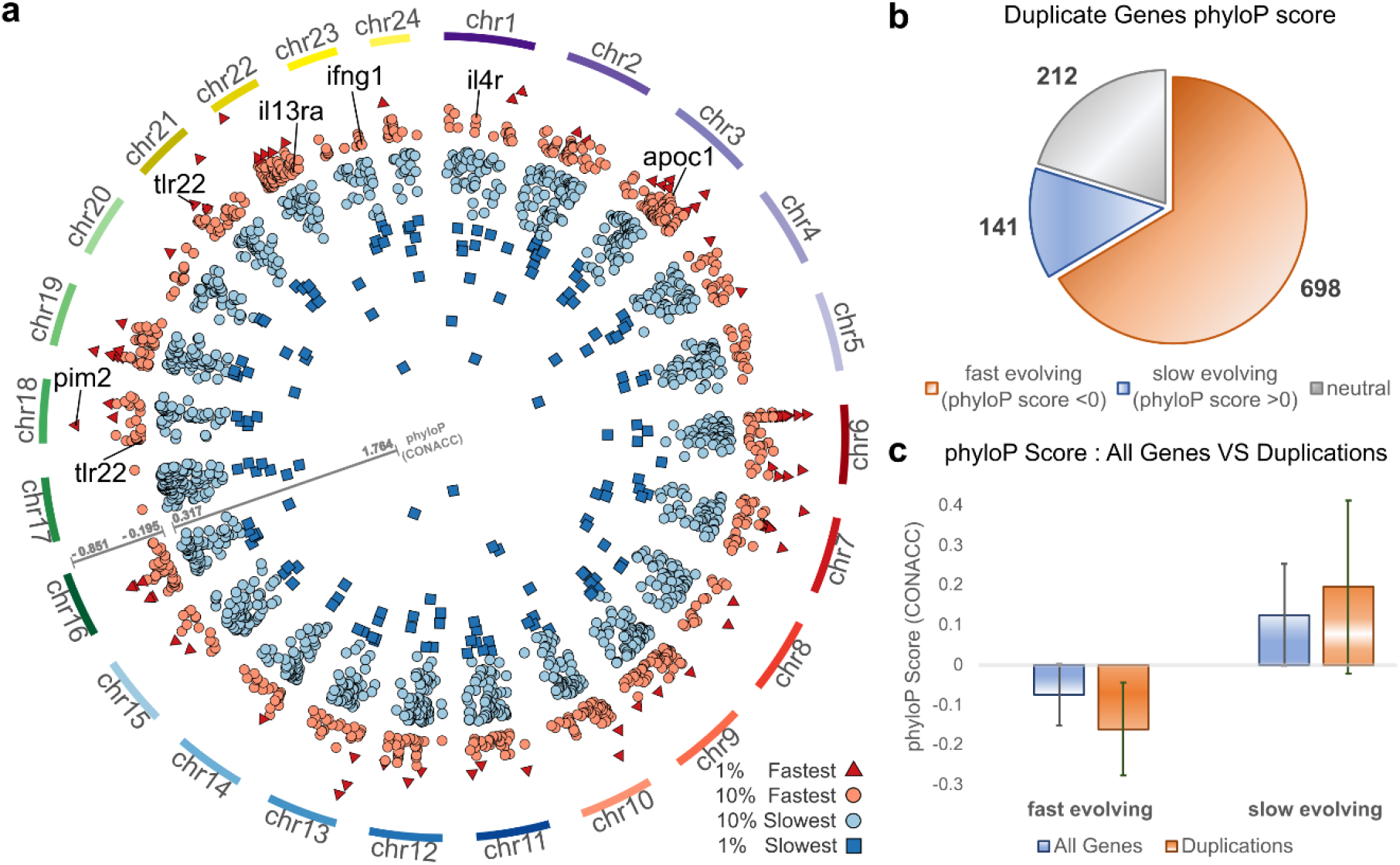
Conservation analysis of meagre gene duplicates. **a**. Circos plot of average phyloP (CONACC) score for top 10% fastest (red) and slowest (blue) evolving genes (red circle: 10% fastest: score <=-0.195, red triangle: 1% fastest: score <=-0.402, blue circle: 10% slowest: score >= 0.317, blue rectangle: 1% slowest: score >= 0.732). **b**. Number of genes from orthogroups with duplications that are fast evolving (phyloP score < 0), slow evolving (phyloP score > 0) or neutrally evolving. **c**. phyloP score of all non-neutrally evolving transcripts and transcripts of genes from orthogroups with duplications, with the average for fast evolving (phyloP score <0) transcripts on the left and slow evolving (phyloP score >0) transcripts on the right.

We then investigated the conservation of 1,051 meagre genes that belong to orthogroups with duplications. This revealed that more than 66.4% (698) of these genes are fast evolving, with only 13.4% (141) characterised as slow evolving and the rest (212) as neutral (Fig.4b). In addition, the average conservation score of fast evolving genes in duplication associated orthogroups was much lower than that of all fast-evolving genes (average score -0.16 VS -0.07, *p*-value = 5.72^-68^), while the score of slow-evolving duplication related genes was higher on average than all slow-evolving genes, but with lower significance (average score 0.19 VS 0.12, *p*-value = 9.38^-5^) (Fig.4c).

The top GO terms associated with fast evolving genes included immune system related functions, leukocyte activation and adhesion, cytokine production, response to external biotic stimulus, largely overlapping with terms enriched in duplication associated orthogroups (Table 3). The orthogroups associating with these functions included MHC2, interleukin, interferon and toll-like receptor genes, including *tlr22, ifng1, il4r* and *il13ra*, which are also among the top 10% fastest evolving genes (Fig.4a). This analysis also revealed enrichment of functions related to lipid/ fatty acid metabolism and specifically to the negative regulation of lipid catabolism (Table 3). Locus *apoc1*, which is associated with these functions, is also found among the top 10% fastest evolving genes (Fig. 4a; phyloP score -0.397). Finally, we also detected fast evolution signatures on cancer related loci *tp53* (phylop score -0.145) and *pim2* (Fig. 4a; phylop score -0.708).

**Table 3.** Fast Evolving Genes Ontology Enrichment. The top table shows selected top terms from GO enrichment analysis of meagre fast evolving genes (phyloP score <0), with certain terms omitted to avoid redundancy. The full table available in Supplementary Table 10. The bottom table shows terms from GO enrichment analysis of meagre fast evolving genes that are associated with lipid metabolism.

|  | term_id | term_name | adjusted_p_value |
| --- | --- | --- | --- |
| Top Terms | GO:0006955 | immune response | 4.61E-67 |
|  | GO:0002684 | positive regulation of immune system process | 3.09E-57 |
|  | GO:0045321 | leukocyte activation | 3.06E-46 |
|  | GO:0006952 | defense response | 5.32E-44 |
|  | GO:0051707 | response to other organism | 2.27E-41 |
|  | GO:0043207 | response to external biotic stimulus | 2.61E-41 |
|  | GO:0002274 | myeloid leukocyte activation | 8.53E-41 |
|  | GO:0002443 | leukocyte mediated immunity | 1.09E-40 |
|  | GO:0001819 | positive regulation of cytokine production | 2.03E-40 |
|  | GO:1903037 | regulation of leukocyte cell-cell adhesion | 3.01E-37 |
| Lipid Metabolism | GO:1901568 | fatty_acid_derivative_metabolic_process | 3.09E-06 |
|  | GO:0050995 | negative_regulation_of_lipid_catabolic_process | 1.05E-05 |
|  | GO:0001676 | long-chain_fatty_acid_metabolic_process | 2.01E-04 |
|  | GO:0035336 | long-chain_fatty-acyl-CoA_metabolic_process | 2.23E-03 |
|  | GO:0050994 | regulation_of_lipid_catabolic_process | 3.02E-03 |
|  | GO:0045833 | negative_regulation_of_lipid_metabolic_process | 1.14E-02 |
|  | GO:0006631 | fatty_acid_metabolic_process | 1.25E-02 |
|  | GO:0016042 | lipid_catabolic_process | 2.50E-02 |
|  | GO:0006638 | neutral_lipid_metabolic_process | 3.09E-02 |
|  | GO:1901570 | fatty_acid_derivative_biosynthetic_process | 4.37E-02 |

### Sciaenid Synteny

To study synteny conservation within Sciaenidae, the genomes of *Larimichthys crocea* and *Collichthys lucidus* were selected for their high contiguity and completeness. First, we investigated conservation of macrosynteny across Sciaenid genomes, filtering orthogroups with a single locus in the meagre and each other member of the family, plotting the position of these single copy orthologs in the 24 chromosomes of each species as a circos plot presented in Figure 5. Synteny is highly conserved within the family, especially between the meagre and *C. lucidus*. However, differences in synteny patterns can be seen, particularly with *L. crocea* chromosomes 1,2,3,4,6 and *C. lucidus* chromosome 1.

**Figure 5.**
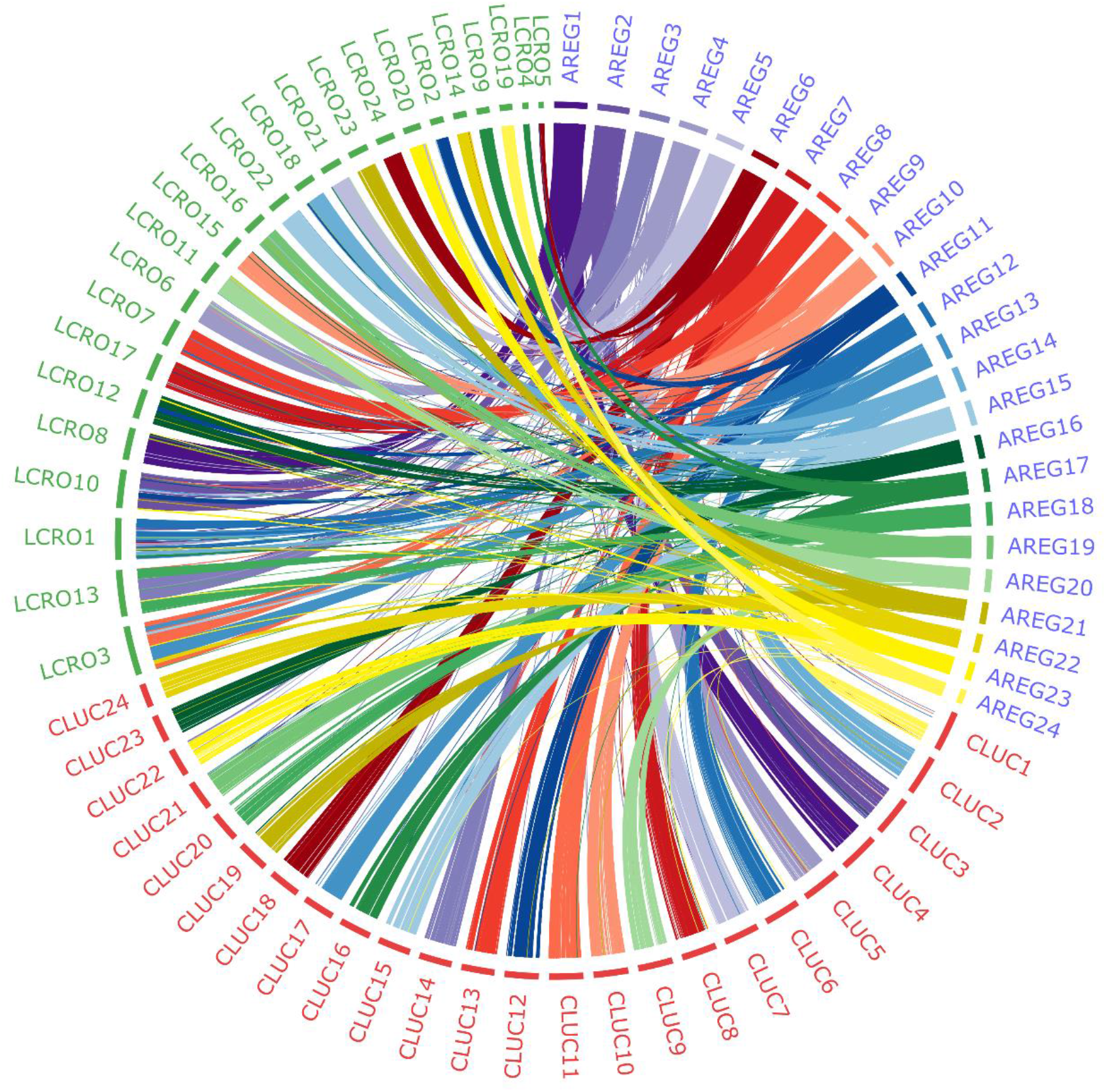
*Sciaenidae* single copy ortholog synteny. Circos plot of one-to-one synteny of single copy orthologous loci in Sciaenidae genomes (CLUC: *Collichthys lucidus*, LCRO: *Larimichthys crocea*, AREG: *Argyrosomus regius* – meagre). Loci in each meagre chromosome (colour coded differently) are connected with coloured lines (corresponding to meagre chromosomal colour codes) to orthologous loci in other genomes (orange for *C. lucidus*, green for *L. crocea*).

### *Sex-related dmrt1* Locus Evolution

Capitalising on the quality of the new meagre assembly and the observed conserved synteny in the family, we studied microsynteny in the *dmrt1* neighbourhood, aiming to obtain insight into the evolution of sex determination in the Sciaenidae family. In this analysis we focused on *dmrt1* and 9 syntenic genes: *hank1a, cfap157, dmrt3, dmrt2a, fam102a, odf2b, rnf183, piwil2* and *golga3*. To obtain a view of the state of the neighbourhood before the Sciaenid ancestor, we used the European sea bass *Dicentrarchus labrax* and the Nile tilapia *Oreochromis niloticus* as outgroups. At the same time, we also searched the recently published genome of *Argyrosomus japonicus*, to better understand the origin of *A. regius* duplications. Both outgroup species exhibited a highly similar structure in the neighbourhood, though a tandem duplication of *piwil2* occurred in *D. labrax* (Fig. 6). Comparison of the four Sciaenidae to these outgroup genomes revealed a series of genomic alterations in the neighbourhood, including translocations, inversions and duplications (Fig. 6). Parsimoniously, the *L. crocea, A. regius* and *A. japonicus* loci suggest an initial translocation of a block comprising *dmrt2a, odf2b and fam102a* and an inversion of *odf2b* and *fam102a* in the Sciaenid ancestor. In *L. crocea* there was a subsequent inversion of *dmrt3* and *dmrt1*. In *C. lucidus*, a segmental block including *dmrt3, cfap157, dmrt1, rnf183, piwil2* duplicated in tandem and was then inverted, with an additional duplication of *cfap157* upstream of these duplicate blocks, while there was an inversion and a translocation of the *dmrt2a, odf2b, fam102a* block downstream of *golga3*, with additional tandem duplications of *fam102a* (3 copies) and *odf2b* (2 copies). In *A. regius*, a series of tandem duplications have resulted in 3 copies of *dmrt3*, 2 copies of *cfap157* and 2 copies of *dmrt1*. Based on the fragmented structure of the second *dmrt1* copy and the short length of the remaining fragments, it is possible that the copy may be pseudogenizing (dmrt1-fr in Fig.6), while comparison to *A. japonicus* suggests that a block duplication of *cfap157* and *dmrt1* happened early in the evolution of the *Argyrosomus* genus. Strikingly, it also reveals that the two tandem duplications of *dmrt3* are very recent and specific to *A. regius*. Phylogenomic reconstruction of the relationships of *dmrt1, dmrt3* and *dmrt2a* (outgroup) proteins for these loci and species also support the duplications of *dmrt1* and *dmrt3* being specific to *C. lucidus* and the *Argyrosomus* genus respectively and not shared. In contrast, the *drmt1* duplication is shared by *A. regius* and *A. japonicus* (Supplementary Fig.2).

**Figure 6.**
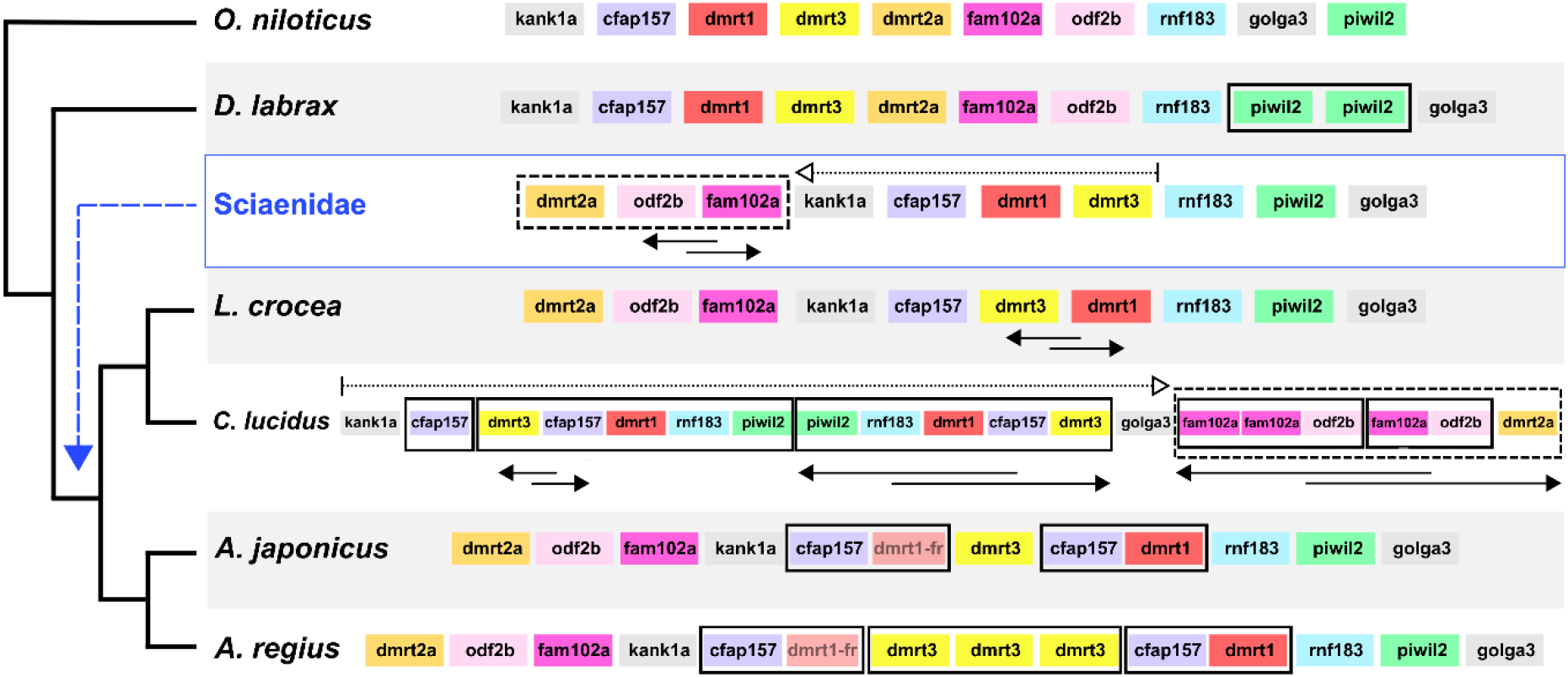
Sciaenidae sex-related *dmrt1* locus synteny and evolution. Synteny of genes in the *dmrt1* neighbourhood. The neighbourhood in each taxon is contained in a large white or grey rectangle, with taxon names on the bottom left. Each gene is represented as a coloured box, with distances and sizes non representative of real quantities. Duplications are indicated by bold black boxes, translocations are indicated by dotted boxes and arrows, inversions of gene blocks are indicated by two-way overlapping black arrows.

## Discussion

### Assembly and Annotation

This work presents the first nuclear genome for the meagre, *Argyrosomus regius*. This new chromosome level genomic assembly has very high completeness, as assessed by two independent methods, kmer counting and conserved ortholog presence/absence. Furthermore, we offer high quality annotation sets for genes, transcripts and repeats, providing the means for further downstream dissection of meagre adaptions and important tools for follow-up experiments on meagre biology. The assembly and scaffolding of the genomic data to chromosomal contiguity was powered by combining our short-long read sequencing strategy with the high-quality linkage map previously constructed for the species. In contrast to other more demanding technologies for sequencing and scaffolding (e.g., Hi-C), our approach achieved optimal results with minimised sequencing effort in a significantly more time and cost-efficient manner.

It is also important to underline how the two-way exchange of information between linkage map and genomic assembly building allowed for the improvement of both resources. Notably, the new genomic assembly was used to assess and validate linkage group structure, in order to subsequently use the linkage groups for scaffolding. As seen in Supplementary Fig.1 and detailed in materials and methods, contig mapping on the linkage map guided the breakage of large LG I into two separate linkage groups, while supporting the discarding of LG XXIV, due to poor and uneven mapping of genomic regions. Following these two amendments, the linkage map successfully guided the scaffolding of the genome to a chromosomal level, with 92.85% of genome length in the 24 chromosomes.

The genome size (696Mb) suggested by the new assembly is slightly larger than that of the large yellow croaker *L. crocea* (679Mb) and smaller than that of *C. lucidus* (877Mb) and the repeat content (25.86%) also ranges between that of the former (18.1%) and the latter (34,68%). The number of annotated genes (24,589) is closer to the number predicted in *L. crocea* (25,401), while *C. lucidus* has more predicted genes than either (28,602). Interestingly, we identified a much larger number of single-copy orthologs between the meagre and *L. crocea* (11,225 genes), than *C. lucidus* (8,175).

### Meagre adaptations highlighted by genomic analyses

The reconstructed phylogeny built from the assigned 2,104 single copy ortholog proteins confirmed the placement of the meagre and Sciaenidae in the expected positions, with *L. crocea* and *C. lucidus* grouping together and the European sea bass placed as the outgroup to the family. Based on the branch length (number of substitutions per site), Sciaenidae genomes appear to be more slowly evolving, on average, compared to other teleost species included, with the meagre genome evolving slower than *L. crocea* and *C. lucidus*.

The short branch of meagre prompted us to look for signatures of selection more closely. For this purpose, we coupled gene duplication search with base-wise conservation analysis both in duplicates and genome-wide, to identify patterns indicating adaptive forces at play. First, ontology analysis of orthogroups with duplications revealed enrichment in many immune system related processes. However, such duplication patterns can be common, because of the continuous interaction of the immune system with the environment of a species. Therefore, we used a strict, comparative approach to filter and isolate the terms specifically associated with meagre duplications. After shortlisting orthogroups with two or more duplications in CAFE that GeneRax also confirms, we selected ontology terms significantly enriched in the meagre and a maximum of three other species. While this may have discarded processes that correspond to potentially real meagre adaptations, we strived to offer high confidence candidates for targeted follow up analyses. Among top biological functions shortlisted in this fashion, categories with potential for such focused studies include cell-cell adhesion (leukocyte cell-cell adhesion, cell-cell adhesion via plasma-membrane adhesion molecules), immune signalling cascade (immune response-activating signal transduction, interferon-gamma-mediated signaling pathway, negative regulation of protein activation cascade) and antigen sampling in mucosal-associated lymphoid tissue.

Complementing the gene duplication data, enrichment analysis carried out on fast evolving genes ordered by conservation score provides further support on meagre adaptations relating to these and other immune system functions, with top enrichment terms in this analysis including leukocyte activation, positive regulation of cytokine production and leukocyte cell-cell adhesion. Interestingly, though not unexpectedly, the majority of genes in orthogroups with duplications exhibit lower than average conservation score. The overlap between these datasets may contribute to the shared enrichment terms observed, though the list of fast evolving genes is much larger than the duplication containing orthogroups. Noteworthy fast evolving immune related genes include interferons and their receptors (*ifng1, ifngr2, igfngr1, igfngr2*), interleukins and their receptors (e.g. *il6st, il6r, il17a,il17c, il17rc, il16, il15l, il15ra, il12ba, il12bb, il12rb2, il10, il10ra, il10rb*), toll-like receptors (*tlr1, tlr5a, tlr7, tlr22*) and major histocompatibility complex class genes, with *mhc2* genes (classified as *mhc2da*) showing both rapid expansion and fast evolution.

In addition to immune adaptations, conservation analysis highlighted fast evolving loci that offer intriguing candidates for studying the evolution and regulation of fast growth and large body size in the meagre. Ontology enrichment revealed fast evolution in genes associated with lipid metabolism, catabolism and fatty acid derivative biosynthesis. This list includes gene candidates with described functions in these processes, such as *apoc1*(Fuior & Gafencu, 2019)(also among the top 10% fastest evolving genes), *bscl2*(Chen et al., 2013), *ces3*(Wei et al., 2010), *dbi*(Bouyakdan et al., 2015) and *fads2*(Glaser, Heinrich, & Koletzko, 2010). This evolutionary signature could indicate adaptations on lipid catabolism, potentially connected to rapid growth, making these loci of great importance for studies on increasing aquaculture efficiency for the species. Intriguingly, we also detected low conservation score on two cancer related loci, tumour protein P53 (*tp53)* and *pim2*. These genes are involved in the regulation of cell cycle, cell death and proliferation(Amir et al., 2017; Kronschnabl, Grünweller, Hartmann, Aigner, & Weirauch, 2020; Thomas, Kelly, & Strasser, 2022; Yan et al., 2003), with *tp53* duplications in elephants previously suggested to be important for countering cancer increase due to large size in these animals(Sulak et al., 2016). Importantly, *pim2* has the fourth fastest evolving transcript in the meagre genome, indicating high selection pressure on the locus. Therefore, we speculate the gene could be involved in promoting cell survival, potentially an adaptation connected to the large body size of the meagre.

### Sciaenid sex related *dmrt1* neighbourhood evolution

Single-copy ortholog synteny between the meagre and the other two Sciaenidae is highly conserved, as seen in Fig.3. Strikingly, although *C. lucidus* is more closely related to *L. crocea*, we observe higher synteny conservation between *C. lucidus* and *A. regius*, with one-to-one chromosome correspondence between all chromosomes and only few small-scale translocations in *C. lucidus* compared to the meagre. In contrast, larger scale rearrangements in *L. crocea* include a fusion of regions in chromosome 1 corresponding to areas from meagre chromosomes 9, 13 and 22, as well as a fusion of regions in chromosome 2 corresponding to meagre chromosomes 3 and 18, with some sizable rearrangements also found in *L. crocea* chromosomes 3, 4 and 6.

Previous studies have suggested Sciaenidae share a relatively stable karyotype of 2n=48(Cai et al., 2019; H. Lin et al., 2021), which is observed in *L. crocea*(Ao et al., 2015), in our new meagre genome (built from a female individual) and previous studies on the meagre(Soares et al., 2012), as well as in the recent *Argyrosomus japonicus* genome(Zhao et al., 2021), which is built from a male individual. Building on this work, we hypothesised that *C. lucidus* may consist a divergent case, with *L. crocea* showing a more typical karyotype of the family. Previous work has suggested *L. crocea* has an XY sex determination system, indicating chromosome 3 as the putative sex chromosome and highlighting *dmrt1* as a sex determination locus candidate(H. Lin et al., 2021). This DM-domain containing transcription factor is a major regulator of male gonad development(Lindeman et al., 2015; Matson et al., 2011) and plays a key role in sex determination in mammals(Matson et al., 2011), birds(Ioannidis et al., 2021), turtles(Ge et al., 2017), frogs(Ma, Rodrigues, Sermier, Brelsford, & Perrin, 2016; Yoshimoto et al., 2010) and other teleost species, including medaka(Lutfalla, Roest Crollius, Brunet, Laudet, & Robinson-Rechavi, 2003; Nanda et al., 2002), zebrafish(Webster et al., 2017), Chinese tongue sole(Cui et al., 2017), spotted scat(Mustapha et al., 2018) and mandarin fish(C. Han et al., 2021).

Our analysis into synteny conservation of *dmrt1* and neighbouring genes revealed these 10 loci have remained in a single highly conserved block across teleost fish (Fig.6; Supplementary Fig.3), with all 10 genes linked in 7 non-Sciaenid teleosts, 9 of the genes retaining synteny in zebrafish and 4 other teleosts and 8 found in human chromosome 9 in two syntenic blocks (Supplementary Fig.3). Comparison to Nile tilapia and European sea bass indicated high synteny conservation in the neighbourhood outside Sciaenidae. As such, the translocation of the *dmrt2a, odf2b, fam102a* block in the Sciaenid ancestor is notable and we suggest the disconnect of *dmrt3* and *dmrt2a* may reflect a change in the regulation of the neighbourhood that would be worth investigating in future studies.

Among Sciaenids, the *L. crocea* area is probably the most similar to the ancestral state, with an inversion of *dmrt3-dmrt1*, while *C. lucidus* presents an exception studied so far with copy number and synteny changes in the area, combining inversion and translocation events with segmental and tandem duplications. As mentioned, *L. crocea* uses an XY sex determination system and *dmrt1* has been highlighted as the sex determination locus, with specific deletions in the neighbourhood found in male fish(A. Lin et al., 2017; H. Lin et al., 2021). In contrast, *C. lucidus* has a complex X_1_X_1_X_2_X_2_/X_1_X_2_Y system, with chromosomes 1 and 7 suggested as the X_1_ and X_2_ chromosomes, while the *dmrt1* neighbourhood is located on chromosome 11(Xiao et al., 2020). The evolution of *C. lucidus* sex determination must have happened in a relatively short time, after the split from the shared ancestor with *L. crocea* circa 14.5 million years ago. Thus, the comparably fast and extensive changes in the *C. lucidus dmrt1* area may have accompanied the transition to this new system, maybe due to a disconnection of *dmrt1* from its ancestral role in male sex determination. Alternatively, the highly modified *dmrt1* neighbourhood could have unknown roles in sex determination or gonad differentiation that remain to be determined.

The structure of the *dmrt1* neighbourhood in the *Argyrosomus* genus is similar to *L. crocea* and the ancestral state. Based on the fragmented, yet detectable sequence of the *dmrt1-fr* copy, we suggest the segmental duplication of *cfap157* and *dmrt1* to be relatively recent in the genus, while the two tandem duplications of *dmrt3* appear specific to *A. regius*, based on the comparison to *A. japonicus*. Considering the roles of *dmrt1* in sex determination and the functions of neighbouring genes in gonadal fate determination and spermatogenesis, these rapid changes in the neighbourhood are highly suggestive of adaptive evolution in an evolutionarily short timeframe. Therefore, we hypothesise the *dmrt1* neighbourhood participates in meagre sex determination, comparable to *L. crocea*, while recent duplications possibly reflect recent evolutionary changes in gonadal differentiation or sex determination in the species.

## Acknowledgements

This study has received funding from the Hellenic Republic and the EU through the “MeagreGen” project under the call “Special Actions – AQUACULTURE” in the Operational Programme “Competitiveness Entrepreneurship and Innovation (EPAnEK 2014-2020)”. The computational work in this study was carried out through the resources of the IMBBC (Institute of Marine Biology, Biotechnology and Aquaculture) HPC facility of the HCMR (Hellenic Centre for Marine Research). Funding for establishing the IMBBC HPC has been received by the MARBIGEN (EU Regpot) project, LifeWatchGreece RI and the CMBR (Centre for the study and sustainable exploitation of Marine Biological Resources) RI.

## Data Accessibility

Raw read data and the meagre assembly have been deposited to the European Nucleotide Archive (ENA) under study accession PRJEB56176. Gene and repeat annotations, functional annotation and gene ontology reference information can be found in the supplementary data files.

## Author Contributions

VP carried out all computational work including genome assembly construction, gene and repeat annotation, phylogenomics analyses, whole genome alignments, duplication analyses, evolutionary analyses, synteny analysis and ontology enrichment and wrote the initial paper draft. CT and TM conceived and designed the experiments and sequencing strategy and supervised the work. JBK and AT performed dissections and nucleic acid extractions. JBK carried out MinION sequencing in IMBBC, HCMR. ON constructed and provided the meagre ddRAD linkage map and assisted with linkage map based scaffolding. CCM, CB, DC contributed to the interpretation of the findings. All authors contributed towards paper writing.

## References

Amir, H., Touboul, T., Sabatini, K., Chhabra, D., Garitaonandia, I., Loring, J. F., … Laurent, L. C. (2017). Spontaneous Single-Copy Loss of TP53 in Human Embryonic Stem Cells Markedly Increases Cell Proliferation and Survival. Stem Cells, 35(4), 872–885. doi: 10.1002/stem.2550

Angelova, N., Danis, T., Lagnel, J., & Tsigenopoulos, C. S. (n.d.). SnakeCube : containerized and automated next-generation sequencing (NGS) pipelines for genome analyses in HPC environments. (3), 2–5. doi: 10.5281/zenodo.4670966

Ao, J., Mu, Y., Xiang, L.-X., Fan, D., Feng, M., Zhang, S., … Chen, X. (2015). Genome Sequencing of the Perciform Fish Larimichthys crocea Provides Insights into Molecular and Genetic Mechanisms of Stress Adaptation. PLOS Genetics, 11(4), e1005118. doi: 10.1371/journal.pgen.1005118

Armstrong, J., Hickey, G., Diekhans, M., Fiddes, I. T., Novak, A. M., Deran, A., … Paten, B. (2020). Progressive Cactus is a multiple-genome aligner for the thousand-genome era. Nature, 587(7833), 246–251. doi: 10.1038/s41586-020-2871-y

Bao, W., Kojima, K. K., & Kohany, O. (2015). Repbase Update, a database of repetitive elements in eukaryotic genomes. Mobile DNA, 6(1), 11. doi: 10.1186/s13100-015-0041-9

Bolger, A. M., Lohse, M., & Usadel, B. (2014). Trimmomatic: a flexible trimmer for Illumina sequence data. Bioinformatics, 30(15), 2114–2120. doi: 10.1093/bioinformatics/btu170

Bouyakdan, K., Taïb, B., Budry, L., Zhao, S., Rodaros, D., Neess, D., … Alquier, T. (2015). A novel role for central ACBP/DBI as a regulator of long-chain fatty acid metabolism in astrocytes. Journal of Neurochemistry, 133(2), 253–265. doi: 10.1111/jnc.13035

Buchfink, B., Reuter, K., & Drost, H.-G. (2021). Sensitive protein alignments at tree-of-life scale using DIAMOND. Nature Methods, 18(4), 366–368. doi: 10.1038/s41592-021-01101-x

Cai, M., Zou, Y., Xiao, S., Li, W., Han, Z., Han, F., … Wang, Z. (2019). Chromosome assembly of Collichthys lucidus, a fish of Sciaenidae with a multiple sex chromosome system. Scientific Data, 6(1). doi: 10.1038/s41597-019-0139-x

Camacho, C., Coulouris, G., Avagyan, V., Ma, N., Papadopoulos, J., Bealer, K., & Madden, T. L. (2009). BLAST+: architecture and applications. BMC Bioinformatics, 10(1), 1–9. doi: 10.1186/1471-2105-10-421

Capella-Gutierrez, S., Silla-Martinez, J. M., & Gabaldon, T. (2009). trimAl: a tool for automated alignment trimming in large-scale phylogenetic analyses. Bioinformatics, 25(15), 1972–1973. doi: 10.1093/bioinformatics/btp348

Chatzifotis, S., Panagiotidou, M., & Divanach, P. (2012). Effect of protein and lipid dietary levels on the growth of juvenile meagre (Argyrosomus regius). Aquaculture International, 20(1), 91–98. doi: 10.1007/s10499-011-9443-y

Chatzifotis, S., Panagiotidou, M., Papaioannou, N., Pavlidis, M., Nengas, I., & Mylonas, C. C. (2010). Effect of dietary lipid levels on growth, feed utilization, body composition and serum metabolites of meagre (Argyrosomus regius) juveniles. Aquaculture, 307(1–2), 65–70. doi: 10.1016/j.aquaculture.2010.07.002

Chen, W., Zhou, H., Liu, S., Fhaner, C. J., Gross, B. C., Lydic, T. A., & Reid, G. E. (2013). Altered lipid metabolism in residual white adipose tissues of Bscl2 deficient mice. PLoS ONE, 8(12). doi: 10.1371/journal.pone.0082526

Cui, Z., Liu, Y., Wang, W., Wang, Q., Zhang, N., Lin, F., … Chen, S. (2017). Genome editing reveals dmrt1 as an essential male sex-determining gene in Chinese tongue sole (Cynoglossus semilaevis). Scientific Reports, 7. doi: 10.1038/srep42213

Danis, T., Papadogiannis, V., Tsakogiannis, A., Kristoffersen, J. B., Golani, D., Tsaparis, D., … Manousaki, T. (2021). Genome Analysis of Lagocephalus sceleratus: Unraveling the Genomic Landscape of a Successful Invader. Frontiers in Genetics, 12. doi: 10.3389/fgene.2021.790850

Darriba, D., Posada, D., Kozlov, A. M., Stamatakis, A., Morel, B., & Flouri, T. (2020). ModelTest-NG: A New and Scalable Tool for the Selection of DNA and Protein Evolutionary Models. Molecular Biology and Evolution, 37(1), 291–294. doi: 10.1093/molbev/msz189

de Coster, W., D’Hert, S., Schultz, D. T., Cruts, M., & van Broeckhoven, C. (2018). NanoPack: visualizing and processing long-read sequencing data. Bioinformatics, 34(15), 2666–2669. doi: 10.1093/bioinformatics/bty149

Dobin, A., Davis, C. A., Schlesinger, F., Drenkow, J., Zaleski, C., Jha, S., … Gingeras, T. R. (2013). STAR: ultrafast universal RNA-seq aligner. Bioinformatics, 29(1), 15–21. doi: 10.1093/bioinformatics/bts635

Emms, D. M., & Kelly, S. (2015). OrthoFinder: solving fundamental biases in whole genome comparisons dramatically improves orthogroup inference accuracy. Genome Biology, 16(1), 1–14. doi: 10.1186/s13059-015-0721-2

Emms, D. M., & Kelly, S. (2019). OrthoFinder: Phylogenetic orthology inference for comparative genomics. Genome Biology, 20(1), 1–14. doi: 10.1186/s13059-019-1832-y

FastQC. (2015, June). Retrieved from https://qubeshub.org/resources/fastqc

Fuior, E. v., & Gafencu, A. v. (2019, December 1). Apolipoprotein c1: Its pleiotropic effects in lipid metabolism and beyond. International Journal of Molecular Sciences, Vol. 20. MDPI AG. doi: 10.3390/ijms20235939

Ge, C., Ye, J., Zhang, H., Zhang, Y., Sun, W., Sang, Y., … Qian, G. (2017). Dmrt1 induces the male pathway in a turtle species with temperature-dependent sex determination. Development (Cambridge), 144(12), 2222–2233. doi: 10.1242/dev.152033

Glaser, C., Heinrich, J., & Koletzko, B. (2010, July). Role of FADS1 and FADS2 polymorphisms in polyunsaturated fatty acid metabolism. Metabolism: Clinical and Experimental, Vol. 59, pp. 993–999. doi: 10.1016/j.metabol.2009.10.022

Guerreiro, I., Castro, C., Antunes, B., Coutinho, F., Rangel, F., Couto, A., … Enes, P. (2020). Catching black soldier fly for meagre: Growth, whole-body fatty acid profile and metabolic responses. Aquaculture, 516. doi: 10.1016/j.aquaculture.2019.734613

Gurevich, A., Saveliev, V., Vyahhi, N., & Tesler, G. (2013). QUAST: Quality assessment tool for genome assemblies. Bioinformatics, 29(8), 1072–1075. doi: 10.1093/bioinformatics/btt086

Haas, B. J. (2003). Improving the Arabidopsis genome annotation using maximal transcript alignment assemblies. Nucleic Acids Research, 31(19), 5654–5666. doi: 10.1093/nar/gkg770

Han, C., Wang, C., Ouyang, H., Zhu, Q., Huang, J., Han, L., … Zhang, Y. (2021). Characterization of dmrts and their potential role in gonadal development of mandarin fish (Siniperca chuatsi). Aquaculture Reports, 21. doi: 10.1016/j.aqrep.2021.100802

Han, M. v., Thomas, G. W. C., Lugo-Martinez, J., & Hahn, M. W. (2013). Estimating Gene Gain and Loss Rates in the Presence of Error in Genome Assembly and Annotation Using CAFE 3. Molecular Biology and Evolution, 30(8), 1987–1997. doi: 10.1093/molbev/mst100

Huerta-Cepas, J., Szklarczyk, D., Heller, D., Hernández-Plaza, A., Forslund, S. K., Cook, H., … Bork, P. (2019). EggNOG 5.0: A hierarchical, functionally and phylogenetically annotated orthology resource based on 5090 organisms and 2502 viruses. Nucleic Acids Research, 47(D1), D309–D314. doi: 10.1093/nar/gky1085

Ioannidis, J., Taylor, G., Zhao, D., Liu, L., Idoko-Akoh, A., Gong, D., … Clinton, M. (2021). Primary sex determination in birds depends on DMRT1 dosage, but gonadal sex does not determine adult secondary sex characteristics. Proceedings of the National Academy of Sciences, 118(10). doi: 10.1073/pnas.2020909118

Katoh, K. (2002). MAFFT: a novel method for rapid multiple sequence alignment based on fast Fourier transform. Nucleic Acids Research, 30(14), 3059–3066. doi: 10.1093/nar/gkf436

Kolmogorov, M., Yuan, J., Lin, Y., & Pevzner, P. A. (2019). Assembly of long, error-prone reads using repeat graphs. Nature Biotechnology, 37(5), 540–546. doi: 10.1038/s41587-019-0072-8

Kovaka, S., Zimin, A. v., Pertea, G. M., Razaghi, R., Salzberg, S. L., & Pertea, M. (2019). Transcriptome assembly from long-read RNA-seq alignments with StringTie2. Genome Biology, 20(1), 278. doi: 10.1186/s13059-019-1910-1

Kronschnabl, P., Grünweller, A., Hartmann, R. K., Aigner, A., & Weirauch, U. (2020). Inhibition of PIM2 in liver cancer decreases tumor cell proliferation in vitro and in vivo primarily through the modulation of cell cycle progression. International Journal of Oncology, 56(2), 448–459. doi: 10.3892/ijo.2019.4936

Krzywinski, M., Schein, J., Birol, I., Connors, J., Gascoyne, R., Horsman, D., … Marra, M. A. (2009). Circos: An information aesthetic for comparative genomics. Genome Research, 19(9), 1639–1645. doi: 10.1101/gr.092759.109

Kumar, S., Suleski, M., Craig, J. M., Kasprowicz, A. E., Sanderford, M., Li, M., … Hedges, S. B. (2022). TimeTree 5: An Expanded Resource for Species Divergence Times. Molecular Biology and Evolution, 39(8). doi: 10.1093/molbev/msac174

Lin, A., Xiao, S., Xu, S., Ye, K., Lin, X., Sun, S., & Wang, Z. (2017). Identification of a male-specific DNA marker in the large yellow croaker (Larimichthys crocea). Aquaculture, 480, 116–122. doi: 10.1016/j.aquaculture.2017.08.009

Lin, H., Zhou, Z., Zhao, J., Zhou, T., Bai, H., Ke, Q., … Xu, P. (2021). Genome-Wide Association Study Identifies Genomic Loci of Sex Determination and Gonadosomatic Index Traits in Large Yellow Croaker (Larimichthys crocea). Marine Biotechnology, 23(1), 127–139. doi: 10.1007/s10126-020-10007-2

Lindeman, R. E., Gearhart, M. D., Minkina, A., Krentz, A. D., Bardwell, V. J., & Zarkower, D. (2015). Sexual cell-fate reprogramming in the ovary by DMRT1. Current Biology, 25(6), 764–771. doi: 10.1016/j.cub.2015.01.034

Lutfalla, G., Roest Crollius, H., Brunet, F. G., Laudet, V., & Robinson-Rechavi, M. (2003). Inventing a Sex-Specific Gene: A Conserved Role of DMRT1 in Teleost Fishes Plus a Recent Duplication in the Medaka Oryzias latipes Resulted in DMY. Journal of Molecular Evolution, 57(SUPPL. 1). doi: 10.1007/s00239-003-0021-4

Ma, W. J., Rodrigues, N., Sermier, R., Brelsford, A., & Perrin, N. (2016). Dmrt1 polymorphism covaries with sex-determination patterns in Rana temporaria. Ecology and Evolution, 6(15), 5107–5117. doi: 10.1002/ece3.2209

Manousaki, T., Tsakogiannis, A., Lagnel, J., Kyriakis, D., Duncan, N., Estevez, A., & Tsigenopoulos, C. S. (2018). Muscle and liver transcriptome characterization and genetic marker discovery in the farmed meagre, Argyrosomus regius. Marine Genomics, 39, 39–44. doi: 10.1016/j.margen.2018.01.002

Matson, C. K., Murphy, M. W., Sarver, A. L., Griswold, M. D., Bardwell, V. J., & Zarkower, D. (2011). DMRT1 prevents female reprogramming in the postnatal mammalian testis. Nature, 476(7358), 101–105. doi: 10.1038/nature10239

Mi, H., & Thomas, P. (2009). PANTHER pathway: an ontology-based pathway database coupled with data analysis tools. Methods in Molecular Biology (Clifton, N.J.), 563, 123–140. doi: 10.1007/978-1-60761-175-2_7

Morel, B., Kozlov, A. M., Stamatakis, A., & Szollosi, G. J. (2020). GeneRax: A tool for species-tree-aware maximum likelihood-based gene family tree inference under gene duplication, transfer, and loss. Molecular Biology and Evolution, 37(9), 2763–2774. doi: 10.1093/molbev/msaa141

Mustapha, U. F., Jiang, D. N., Liang, Z. H., Gu, H. T., Yang, W., Chen, H. P., … Li, G. L. (2018). Male-specific Dmrt1 is a candidate sex determination gene in spotted scat (Scatophagus argus). Aquaculture, 495, 351–358. doi: 10.1016/j.aquaculture.2018.06.009

Mylonas, C. C., Mitrizakis, N., Castaldo, C. A., Cerviño, C. P., Papadaki, M., & Sigelaki, I. (2013). Reproduction of hatchery-produced meagre Argyrosomus regius in captivity II. Hormonal induction of spawning and monitoring of spawning kinetics, egg production and egg quality. Aquaculture, 414–415, 318–327. doi: https://doi.org/10.1016/j.aquaculture.2013.09.008

Mylonas, C. C., Mitrizakis, N., Papadaki, M., & Sigelaki, I. (2013). Reproduction of hatchery-produced meagre Argyrosomus regius in captivity I. Description of the annual reproductive cycle. Aquaculture, 414–415, 309–317. doi: https://doi.org/10.1016/j.aquaculture.2013.09.009

Nanda, I., Kondo, M., Hornung, U., Asakawa, S., Winkler, C., Shimizu, A., … Schartl, M. (2002). A duplicated copy of DMRT1 in the sex-determining region of the Y chromosome of the medaka, Oryzias latipes. Proceedings of the National Academy of Sciences, 99(18), 11778–11783. doi: 10.1073/pnas.182314699

Nguyen, L. T., Schmidt, H. A., von Haeseler, A., & Minh, B. Q. (2015). IQ-TREE: A fast and effective stochastic algorithm for estimating maximum-likelihood phylogenies. Molecular Biology and Evolution, 32(1), 268–274. doi: 10.1093/molbev/msu300

Niknafs, Y. S., Pandian, B., Iyer, H. K., Chinnaiyan, A. M., & Iyer, M. K. (2017). TACO produces robust multisample transcriptome assemblies from RNA-seq. Nature Methods, 14(1), 68–70. doi: 10.1038/nmeth.4078

Nousias, O., Oikonomou, S., Manousaki, T., Papadogiannis, V., Angelova, N., Tsaparis, D., … Tsigenopoulos, C. S. (2022). Linkage mapping, comparative genome analysis, and QTL detection for growth in a non-model teleost, the meagre Argyrosomus regius, using ddRAD sequencing. Scientific Reports, 12(1). doi: 10.1038/s41598-022-09289-4

Nousias, Orestis, Tzokas, K., Papaharisis, L., Ekonomaki, K., Chatziplis, D., Batargias, C., & Tsigenopoulos, C. S. (2021). Genetic Variability, Population Structure, and Relatedness Analysis of Meagre Stocks as an Informative Basis for New Breeding Schemes. Fishes, 6(4), 78. doi: 10.3390/fishes6040078

Papadakis, I. E., Kentouri, M., Divanach, P., & Mylonas, C. C. (2013). Ontogeny of the digestive system of meagre Argyrosomus regius reared in a mesocosm, and quantitative changes of lipids in the liver from hatching to juvenile. Aquaculture, 388–391(1), 76–88. doi: 10.1016/j.aquaculture.2013.01.012

Pollard, K. S., Hubisz, M. J., Rosenbloom, K. R., & Siepel, A. (2010). Detection of nonneutral substitution rates on mammalian phylogenies. Genome Research, 20(1), 110–121. doi: 10.1101/gr.097857.109

Ramos-Júdez, S., González, W., Dutto, G., Mylonas, C. C., Fauvel, C., & Duncan, N. (2019). Gamete quality and management for in vitro fertilisation in meagre (Argyrosomus regius). Aquaculture, 509, 227–235. doi: 10.1016/j.aquaculture.2019.05.033

Raudvere, U., Kolberg, L., Kuzmin, I., Arak, T., Adler, P., Peterson, H., & Vilo, J. (2019). G:Profiler: A web server for functional enrichment analysis and conversions of gene lists (2019 update). Nucleic Acids Research, 47(W1), W191–W198. doi: 10.1093/nar/gkz369

Rhie, A., Walenz, B. P., Koren, S., & Phillippy, A. M. (2020). Merqury: Reference-free quality, completeness, and phasing assessment for genome assemblies. Genome Biology, 21(1), 1–29. doi: 10.1186/s13059-020-02134-9

Ribeiro, L., Moura, J., Santos, M., Colen, R., Rodrigues, V., Bandarra, N., … Dias, J. (2015, October 1). Effect of vegetable based diets on growth, intestinal morphology, activity of intestinal enzymes and haematological stress indicators in meagre (Argyrosomus regius). Aquaculture, Vol. 447, pp. 116–128. Elsevier B.V. doi: 10.1016/j.aquaculture.2014.12.017

Siepel, A., & Haussler, D. (2004). Phylogenetic Estimation of Context-Dependent Substitution Rates by Maximum Likelihood. Molecular Biology and Evolution, 21(3), 468–488. doi: 10.1093/molbev/msh039

Simão, F. A., Waterhouse, R. M., Ioannidis, P., Kriventseva, E. v., & Zdobnov, E. M. (2015). BUSCO: Assessing genome assembly and annotation completeness with single-copy orthologs. Bioinformatics, 31(19), 3210–3212. doi: 10.1093/bioinformatics/btv351

Smit, A. H. R. & G. P. (2015). RepeatMasker Open-4.0. 2013-2015 <http://www.repeatmasker.org>..

Soares, F., Leitão, A., Moreira, M., de Sousa, J., Almeida, A., Barata, M., … Ribeiro, L. (2012). Sarcoma in the thymus of juvenile meagre Argyrosomus regius reared in an intensive system. Diseases of Aquatic Organisms, 102(2), 119–127. doi: 10.3354/dao02545

Stamatakis, a, Ludwig, T., & Meier, H. (2003). RAxML: A Parallel Program for Phylogenetic Tree Inference. Poster Abstract in Proceedings of 2nd European Conference on Computational Biology (ECCB2003), 325–326.

Stamatakis, A. (2014). RAxML version 8: a tool for phylogenetic analysis and post-analysis of large phylogenies. Bioinformatics, 30(9), 1312–1313. doi: 10.1093/bioinformatics/btu033

Stanke, M., & Morgenstern, B. (2005). AUGUSTUS: A web server for gene prediction in eukaryotes that allows user-defined constraints. Nucleic Acids Research, 33(SUPPL. 2), 465–467. doi: 10.1093/nar/gki458

Sulak, M., Fong, L., Mika, K., Chigurupati, S., Yon, L., Mongan, N. P., … Lynch, V. J. (2016). TP53 copy number expansion is associated with the evolution of increased body size and an enhanced DNA damage response in elephants. doi: 10.7554/eLife.11994.001

Tang, H., Zhang, X., Miao, C., Zhang, J., Ming, R., Schnable, J. C., … Lu, J. (2015). ALLMAPS: robust scaffold ordering based on multiple maps. Genome Biology, 16(1), 3. doi: 10.1186/s13059-014-0573-1

Thomas, A. F., Kelly, G. L., & Strasser, A. (2022, May 1). Of the many cellular responses activated by TP53, which ones are critical for tumour suppression? Cell Death and Differentiation, Vol. 29, pp. 961–971. Springer Nature. doi: 10.1038/s41418-022-00996-z

Vallecillos, A., María-Dolores, E., Villa, J., Rueda, F. M., Carrillo, J., Ramis, G., … Armero, E. (2022). Development of the First Microsatellite Multiplex PCR Panel for Meagre (Argyrosomus regius), a Commercial Aquaculture Species. Fishes, 7(3), 117. doi: 10.3390/fishes7030117

Venturini, L., Caim, S., Kaithakottil, G. G., Mapleson, D. L., & Swarbreck, D. (2018). Leveraging multiple transcriptome assembly methods for improved gene structure annotation. GigaScience, 7(8). doi: 10.1093/gigascience/giy093

Vurture, G. W., Sedlazeck, F. J., Nattestad, M., Underwood, C. J., Fang, H., Gurtowski, J., & Schatz, M. C. (2017). GenomeScope: fast reference-free genome profiling from short reads. Bioinformatics, 33(14), 2202–2204. doi: 10.1093/bioinformatics/btx153

Walker, B. J., Abeel, T., Shea, T., Priest, M., Abouelliel, A., Sakthikumar, S., … Earl, A. M. (2014). Pilon: An integrated tool for comprehensive microbial variant detection and genome assembly improvement. PLoS ONE, 9(11). doi: 10.1371/journal.pone.0112963

Webster, K. A., Schach, U., Ordaz, A., Steinfeld, J. S., Draper, B. W., & Siegfried, K. R. (2017). Dmrt1 is necessary for male sexual development in zebrafish. Developmental Biology, 422(1), 33–46. doi: 10.1016/j.ydbio.2016.12.008

Wei, E., ben Ali, Y., Lyon, J., Wang, H., Nelson, R., Dolinsky, V. W., … Lehner, R. (2010). Loss of TGH/Ces3 in Mice Decreases Blood Lipids, Improves Glucose Tolerance, and Increases Energy Expenditure. Cell Metabolism, 11(3), 183–193. doi: 10.1016/j.cmet.2010.02.005

Wong, W. Y., & Simakov, O. (2019). RepeatCraft: a meta-pipeline for repetitive element de-fragmentation and annotation. Bioinformatics, 35(6), 1051–1052. doi: 10.1093/bioinformatics/bty745

Xiao, J., Zou, Y., Xiao, S., Chen, J., Wang, Z., Wang, Y., … Cai, M. (2020). Development of a PCR-based genetic sex identification method in spinyhead croaker (Collichthys lucidus). Aquaculture, 522. doi: 10.1016/j.aquaculture.2020.735130

Xu, T., Li, Y., Chu, Q., & Zheng, W. (2021). A chromosome-level genome assembly of the red drum, Sciaenops ocellatus. Aquaculture and Fisheries, 6(2), 178–185. doi: 10.1016/j.aaf.2020.08.001

Xu, Z., & Wang, H. (2007). LTR_FINDER: an efficient tool for the prediction of full-length LTR retrotransposons. Nucleic Acids Research, 35(Web Server), W265–W268. doi: 10.1093/nar/gkm286

Yan, B., Zemskova, M., Holder, S., Chin, V., Kraft, A., Koskinen, P. J., & Lilly, M. (2003). The PIM-2 Kinase Phosphorylates BAD on Serine 112 and Reverses BAD-induced Cell Death. Journal of Biological Chemistry, 278(46), 45358–45367. doi: 10.1074/jbc.M307933200

Yoshimoto, S., Ikeda, N., Izutsu, Y., Shiba, T., Takamatsu, N., & Ito, M. (2010). Opposite roles of DMRT1 and its W-linked paralogue, DM-W, in sexual dimorphism of Xenopus laevis: Implications of a ZZ/ZW-type sex-determining system. Development, 137(15), 2519–2526. doi: 10.1242/dev.048751

Zafeiropoulos, H., Gioti, A., Ninidakis, S., Potirakis, A., Paragkamian, S., Angelova, N., … Pafilis, E. (2021). 0s and 1s in marine molecular research: a regional HPC perspective. GigaScience, 10(8). doi: 10.1093/gigascience/giab053

Zhao, L., Xu, S., Han, Z., Liu, Q., Ke, W., Liu, A., & Gao, T. (2021). Chromosome-Level Genome Assembly and Annotation of a Sciaenid Fish, Argyrosomus japonicus. Genome Biology and Evolution, 13(2). doi: 10.1093/gbe/evaa246

