## Supplementary Data for "Chromosome assembly for the meagre, *Argyrosomus regius*, reveals species adaptations and sciaenid sex-related locus evolution"

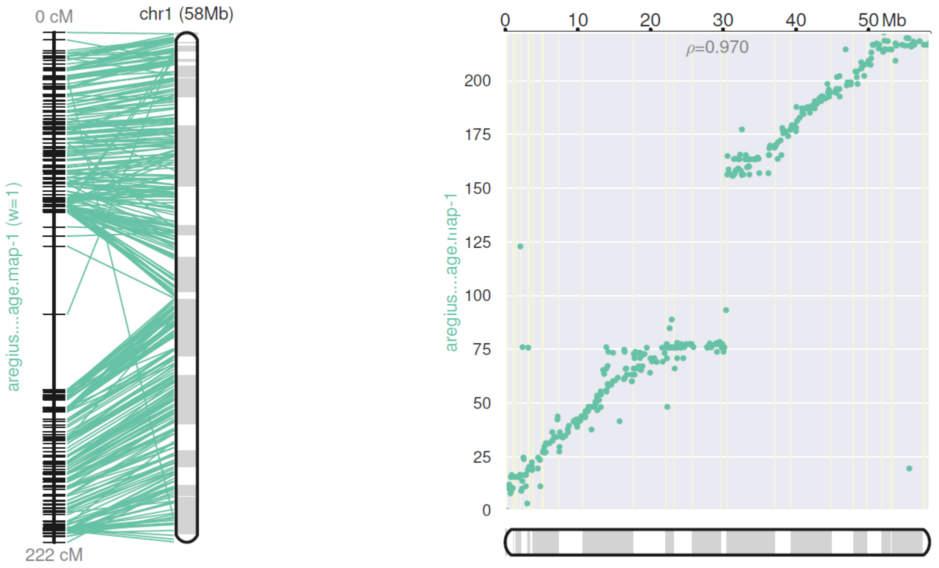


**Supplementary Figure 1. Original Linkage Group I Mapping to Assembly Contigs.** Scaffolding via ALLMAPS revealed assembly contigs mapped to two non-overlapping regions of the original Linkage group I of the meagre linkage map, suggesting the linkage group was a merge of two constituent groups.

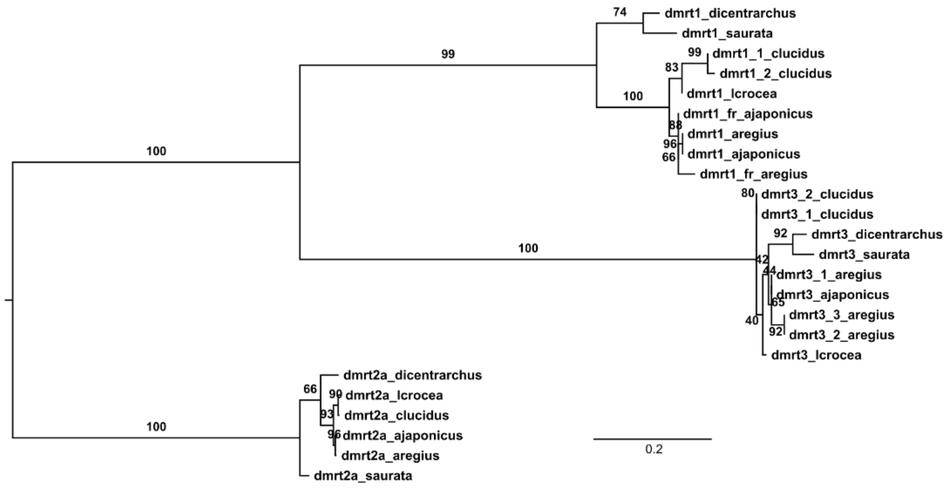


**Supplementary Figure 2. Phylogenomic reconstruction of sciaenid DMRT1, DMRT2 and DMRT3 protein relationships.** DMRT protein sequences from each species were aligned using MAFFT, followed by trimming with trimAl (gap threshold of 20%). Tree inference was carried out with RAxML-ng, using the JTT model (selected by ModelTest-NG) and bootstrap resampling with 100 bootstrap replicates.

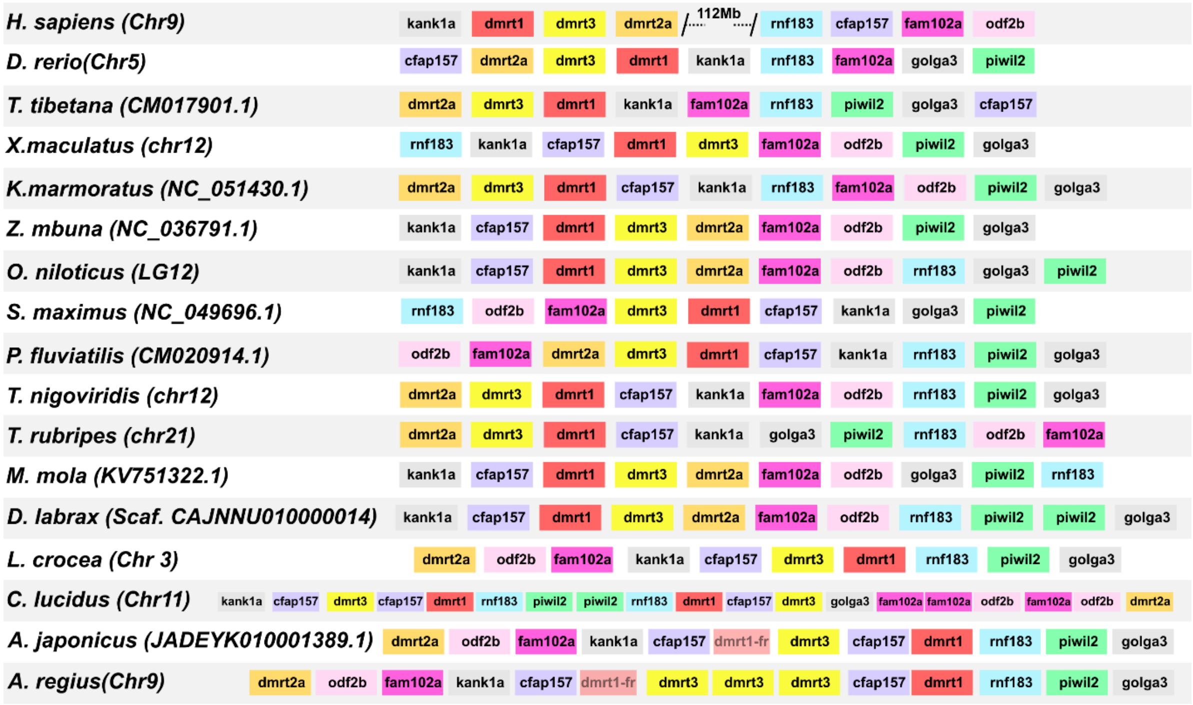


**Supplementary Figure 3. Extended synteny diagram for *dmrt1* neighbourhood genes in different teleosts and human.** The chromosome or scaffold of the neighbourhood in each species is given next to the species name. Genes are colour coded by gene name, following the same code as in main Figure 6.

### Supplementary table 1. Genome sequencing quality statistics.

### Supplementary table 2. Transcriptome sequencing quality statistics.

### Supplementary table 3. ENSEMBL proteomes used for homology evidence during annotation.

### Supplementary table 4. Genomes used for comparative genomics analyses.

**Supplementary table 5. Meagre OrthoFinder Hierarchical OrthoGroups (HOGs) with duplications.** These HOGs were selected through a comparison of CAFE and GeneRax output, as described in Materials and Methods.

**Supplementary table 6. GO enrichment analysis via gprofiler on duplication containing orthogroups from all species.** Ontology terms (GO biological) significantly enriched (adjusted p value <= 0.1) in meagre duplications are found on the top of the list, ranked by the number of other species in which each term is also significantly enriched.

**Supplementary table 7. Fast evolving transcript phyloP (CONACC) scores.**

**Supplementary table 8. Slow evolving transcripts phyloP (CONACC) scores.**

**Supplementary table 9. Analysis of phyloP score for meagre genes in duplication containing orthogroups.**

**Supplementary table 10. GO enrichment analysis via gprofiler on fast evolving genes (phyloP score <0).**

**Supplementary table 11. Orthofinder orthogroups (HOGs) used in the study.**
